## Supplementary figures and images for "Endocytic myosin-1 is a force-insensitive, power-generating motor" (from another version of this preprint: version 2, 2023-05-17)

### Supplemental Figure S1

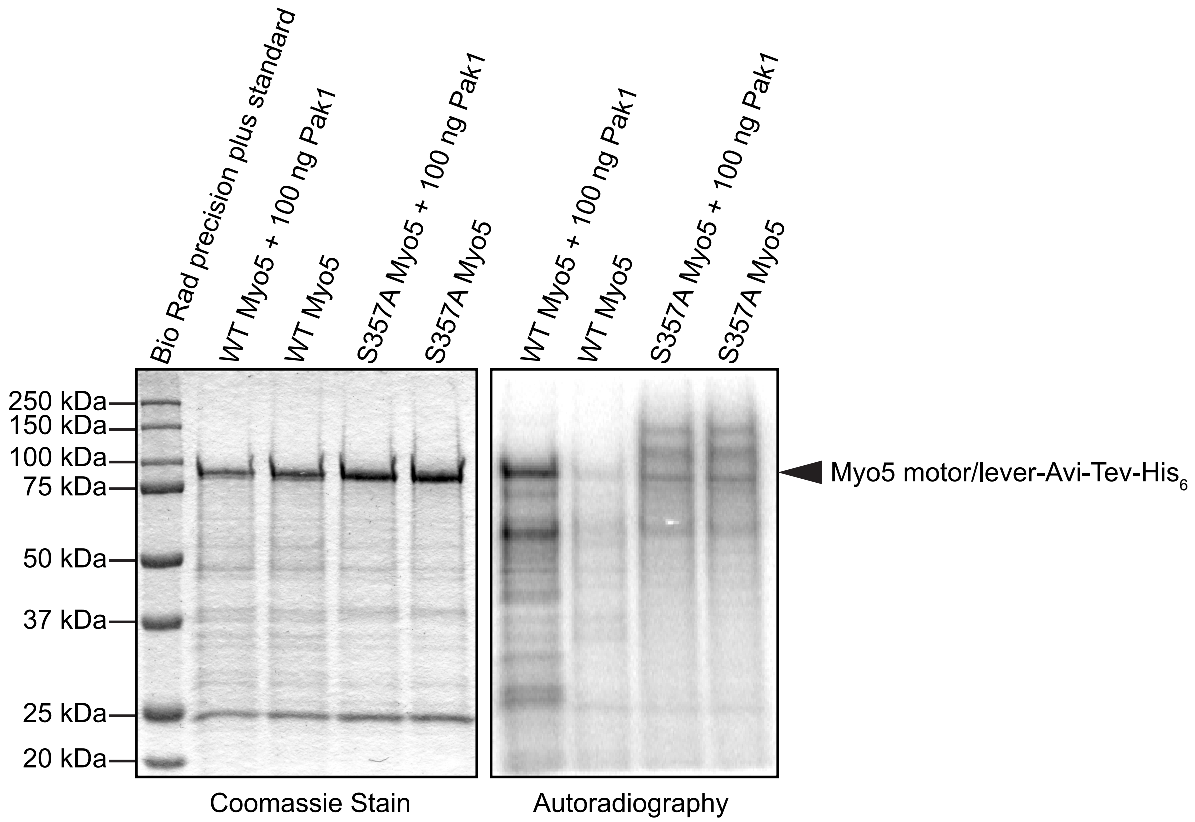
